# Efficient Electric Field Simulations for Transcranial Brain Stimulation

**DOI:** 10.1101/541409

**Authors:** Guilherme B Saturnino, Kristoffer H Madsen, Axel Thielscher

**Author notes:** Shared last authorship (Kristoffer H Madsen, Axel Thielscher).

## Abstract

**Objective:** Transcranial magnetic stimulation (TMS) and transcranial electric stimulation (TES) modulate brain activity non-invasively by generating electric fields either by electromagnetic induction or by injecting currents via skin electrodes. Numerical simulations based on anatomically detailed head models of the TMS and TES electric fields can help us to understand and optimize the spatial stimulation pattern in the brain. However, most realistic simulations are still slow, and their numerical accuracy and the factors that influence it have not been evaluated in detail so far.

**Approach:** We present and validate a new implementation of the Finite Element Method (FEM) for TMS and TES that is based on modern algorithms and libraries. We also evaluate the convergence of the simulations and give estimates for the discretization errors.

**Main results:** Comparisons with analytical solutions for spherical head models validate our new FEM implementation. It is five to ten times faster than previous implementations. The convergence results suggest that accurately capturing the tissue geometry in addition to choosing a sufficiently high mesh density is of fundamental importance for accurate simulations.

**Significance:** The new implementation allows for a substantial increase in computational efficiency of TMS and TES simulations. This is especially relevant for applications such as the systematic assessment of model uncertainty and the optimization of multi-electrode TES montages. The results of our systematic error analysis allow the user to select the best tradeoff between model resolution and simulation speed for a specific application. The new FEM code will be made openly available as a part of our open-source software SimNIBS 3.0.

## 1 Introduction

Transcranial magnetic stimulation (TMS) and transcranial weak electric stimulation (TES) are the two best-established methods for non-invasive transcranial brain stimulation (TBS). Both use electric fields to modulate neural activity in a target brain region or a network of brain regions, but the two methods differ in the mechanism used to generate the field: TMS employs electromagnetic induction, while TES injects currents into the skin via surface electrodes. The ability to transcranially modulate brain activity without serious adverse effects makes TBS a valuable research tool in neuroscience [1,2] and possibly also an effective treatment for several psychiatric and neurological diseases [3]. However, the physiological and behavioral TBS effects are still subject to large inter-subject variability [4,5], which hampers the more wide-spread use of TBS in clinical applications [5].

As TMS and TES use electric fields as mechanism of action to modulate the membrane potential of the neural cells in the brain [3], inter-individual variations of the generated fields is likely a key factor that contributes to the observed physiological and behavioral variability. For both stimulation methods, the electric field distribution is strongly influenced by the anatomical distribution of the head tissues, often in complex and counter-intuitive ways [6–10]. Accurately modeling the electric field distribution generated in the brain, based on individualized models of the head anatomy, is thus important to establish a more stringent control of spatial targeting and dosing. In fact, it has been shown that computational models can help predict stimulation outcome in the case of TMS [9].

There is an increasing interest in tools that perform electric field modelling for TBS [11–13]. However, field simulations with most of the popular packages such as our software SimNIBS (www.simnibs.org) still have high computational cost. A single simulation currently requires several minutes on a standard PC, which is inconvenient for the user. Equally important, this also slows down the development and broad adoption of more advanced applications of realistic and individualized field calculations that rely on the evaluation of many simulations, such as the optimization of multichannel TES montages [14–16] or the systematic uncertainty quantification of the simulation outcome [17–19]. To the best of our knowledge, studies examining the efficiency and accuracy of any of such tools from a numerical perspective are so far still lacking.

In the current work, we present and validate a new implementation of the Finite Element Method (FEM) for TMS and TES that uses modern algorithms and libraries. We first present the basic mathematical equations that underlie the TMS and TES electric fields and discuss how they can be numerically solved using the finite element method (FEM). Afterwards, we describe and validate our new FEM implementation that will be part of a future version of our open source simulation software SimNIBS [11]. We compare the new with the current FEM implementation, demonstrating speed ups of four to nine times and approximately four times decrease in memory requirements, without changes in accuracy or need for special hardware. We conclude with a demonstration of the relative contributions of the FE mesh density and the anatomical fidelity of the modelled tissue boundaries to the overall simulation accuracy.

## 2 Methods

### 2.1 Equations for calculating the TMS and TES electric fields

The equations governing the electric field ***E*** caused by a TMS coil are [20]:

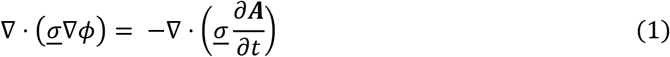

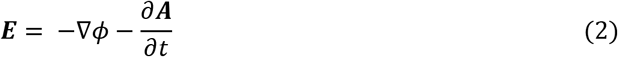

The symbol <u>*σ*</u> denotes the tissue-specific ohmic conductivity, which can be either scalar or a 3×3 symmetric positive definite tensor. In this study, we will assume scalar, piecewise-constant conductivities, with the values varying from 0.01 S/m in bone to around 1.6 S/m in cerebro-spinal fluid (CSF) [6]. ***A*** denotes the magnetic vector potential of the TMS coil, which depends on the coil’s shape, position, and the current flow in the coil wires. The magnetic vector potential (more specifically, its temporal derivative) can be understood as the electric field which the TMS coil would induce in an infinite homogenous conductor. It is also sometimes referred to as the primary electric field. The symbol *ϕ* represents an electrical potential that can be understood as the source of a secondary electric field caused by tissue boundaries or, more generally, by variations in conductivity, partly counteracting the primary field. Additionally, we assume homogeneous Neumann boundary conditions in the entire boundary (that is, there is no current flow to the outside of the head). This model is based on the quasistatic approximation of Maxwell’s equations at low frequencies, which can be safely applied for the TMS simulations [21,22]. This means that, while the current through the TMS coil varies over time, modifying the ***A*** field and by consequence the electric field, we can separate ***E***(***r***, *t*) into two components such that ***E***(***r***, *t*) = ***E***(***r***) ^*∂I*(*t*)/*∂t*^. ***E***(***r***) is a spatial component, calculated using Eq. 2, and *∂I*(*t*)/*∂t* is a temporal component, given solely by the pulse shape.

In the case of TES, the electric potential is governed by a simple Laplace equation [23]:

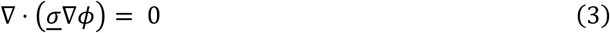

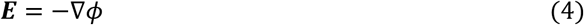

with *ϕ* being the electrical potential caused by the external stimulation, modelled by setting Dirichlet boundary conditions (that is, setting fixed electric potentials) on the electrode surfaces and homogeneous Neumann boundary conditions elsewhere. TES intensity is normally controlled by setting the current flow through the electrodes, and not by setting potential differences. To adjust the simulations to a given current flow, SimNIBS goes through the steps

1. Solve equation 3 setting the potential at an arbitrary reference electrode to 0 V and at an arbitrary active electrode surface to 1V.
2. With the solution, calculate the current flow through the cathode and through the anode.
3. Correct the solution such that the current flow matches the set value. This can be done by a simple linear scaling.
4. Repeat in the case 3 or more electrodes are present

It is also possible to set-up the simulation using only Neumann boundary conditions, in which case the current calibration step would not be necessary. However, this assumes that the current flux through the entire electrode surface is constant, which might be a wrong assumption specially in large electrodes or when connecting many electrodes to a single stimulator channel. However, setting simulations with Neumann boundary conditions can be advantageous when dealing with many small and independently controlled electrodes or when building leadfield matrices for TES optimization [14].

When applying TES with alternating currents, the quasistatic approximation also applies [24]. This means that the electric field at any instant can be calculated by only taking into consideration the current flow in the electrodes at that instant. Additionally, the electric potential is linear with respect to the input currents. This means that we do not need to run a new simulation for each time point, and can instead scale and sum simulation results in the right proportions such that we obtain the desired currents through the electrodes and by consequence the resulting electric field.

Further information about the differential equations governing the TMS and TES electric fields can be found in the Supplementary Material.

### 2.2 Finite Element Method for Electric Field Simulations

Equation 1 and 3, which give us the electric potentials for TMS and TES, respectively, have analytical solutions only in very simple geometries such as spheres [25]. To obtain the electric fields in a realistic head model, we must resort to numerical methods. Here, we apply the Finite Element Method (FEM) [26], a well-established numerical method for obtaining approximate solutions to partial differential equations.

We chose FEM because it offers an elegant and efficient framework to model complex geometries such as the human head. This is done first by discretizing the domain (such as the head) into small sections with simple geometric shape called *elements*. Here, we use tetrahedral elements, but usage of other shapes such as hexahedra is also possible. The elements do not overlap and share their vertices (or *nodes*) with several other elements. The discretized domain, defined by its nodes and elements is called a *mesh*. This simple approach is very effective in representing complicated geometries, such as the sulci and gyri in the human brain. FEM requires the choice of the type of functions (termed *basis functions*) that are used to model the spatial variations of the solution (i.e, the electric potential) within the domain. Here, we use nodal linear basis functions, i.e. we define one basis function per node per element. The basis function has a value of one in its corresponding node and decays linearly within each element, reaching a value of zero in the other element nodes. Outside the element, the basis function has a value of zero. This makes the problem computationally efficient as only immediate neighbors needs to be evaluated meaning that computations can be formulated with sparse matrices.. Finally, we transform Equations 1 and 3 into their weak forms and use the Galerkin method to derive a system of equations based on the linear basis functions.

The steps above, described in detail in the Supplementary Material, results in a system of equations of the form

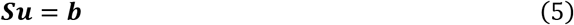

where ***S*** is a sparse matrix of the size *N*_*dof*_ × *N*_*dof*_ and is also denoted as stiffness matrix. *N*_*dof*_ denotes the number of degrees of freedom, which roughly correspond to the number of nodes in the mesh in our case. ***u*** is the electric potential at the nodes and the right-hand side ***b*** contains information about boundary conditions and the *d****A***/*dt* field in the TMS case.

By solving the TMS and TES systems, we obtain the electric potential. As the quantity of interest is the electric field ***E***, we need to compute the negative gradient of *ϕ* to obtain ***E*** in case of TES (eq. 4). For TMS, we have to sum the negative gradient of *ϕ* with the negative of the temporal derivative of the magnetic vector potential to obtain ***E*** (eq. 2). In our FEM implementation that uses linear basis functions for the electric potentials, the results from the gradient calculations will be piecewise-constant. That is, the electric field has a constant value within each element and is discontinuous across element boundaries. Because of these discontinuities, directly using the piecewise-constant gradients to get the generated electric fields will decrease the numerical accuracy of the simulations. To improve on the original solutions, we use the superconvergent patch recovery (SPR) procedure [27] to recover nodal values for the electric fields, and in turn to obtain more accurate interpolations of the electric field values at any position in the head mesh. However, as the electric field is discontinuous across tissue boundaries due to the abrupt changes in conductivity, we calculate the recovered values for each tissue separately.

Calculating the electric field using FEM starts by assembling the stiffness matrix ***S*** and the right-hand side ***b*** from Equation 5. We implemented a computationally efficient vectorized algorithm for stiffness matrix assembly [28] in Python 3, using the NumPy and SciPy packages for efficient vector computations. This implementation vectorizes the element-wise *for loop* usually applied in the assembly of stiffness matrices, allowing for large increases in computational efficiency when using computational libraries that support vector operations.

Afterwards, we need to *solve* the system in Equation 5. ***S*** is a large, sparse matrix. As the size of ***S*** precludes its direct inversion, FEM employs solvers that can obtain the solution ***u*** without explicitly building ***S***^−1^. Many such solvers, and especially the most efficient ones, require positive definite matrices. The matrix ***S*** is already positive definite in the case of TES. However, in the TMS case, it is positive semidefinite. We can however make it positive definite by setting a Dirichlet boundary condition on a single node to “ground” the model, thereby eliminating the floating potential. This does not modify the resulting electric field, as also confirmed in the validation results shown below.

Solvers for sparse systems can be classified as either *direct solvers* or *iterative solvers*. Direct solvers will typically factor the matrix ***S*** into its LU (lower/upper triangular) form [29]. Considering the typical matrix size for the problem at hand (> 10^5^ × 10^5^), direct solvers suffer from large time and memory requirements on forming and storing the factorized form of the matrix. For our implementation, we therefore resort to iterative solvers that to not need to fully decompose the matrix and therefore have lower memory footprints. Iterative solvers start with a guess solution and iteratively reduce the error in the solution. For positive definite matrices, the conjugate gradient (CG) method is a highly efficient iterative method [29]. Iterative methods can achieve a massive speedup by usage of a *preconditioner* [29]. In a simple way, a preconditioner approximates the solution of the linear system and by that improves the efficiency of the subsequent iterative solver. Recent developments have resulted in new and highly efficient preconditioners, such as algebraic multigrid preconditioners [30,31]. In our implementation, we combine an effective iterative solver and a modern preconditioner to achieve very fast and memory efficient FEM calculations. For that, SimNIBS 3.0 interfaces directly with PETSc [32], a powerful scientific computing library which offers a homogeneous interface to a large set of solvers and preconditioners with a very low overhead. Specifically, SimNIBS 3.0 solves the FEM system by default using the preconditioned CG method, with a relative error of 10^−10^ and with the *BommerAMG* preconditioner from *hypre* [33].

### 2.3 Validation in a Spherical Phantom

To validate our FEM implementation, we used a geometric model which consists of five concentric spherical shells of radii 75, 78, 80, 86 and 92 mm [34], shown in Figure 1. The shells emulate the outer boundaries of white matter (WM), gray matter (GM), cerebro-spinal fluid (CSF), skull and skin, respectively. Such a simple model allows us to calculate solutions analytically, and therefore directly evaluate numerical accuracy. We used six different resolutions for the sphere phantoms, shown in Table 1. The FEM solutions were compared to the analytical results by evaluating the electric fields at the nodes of a sphere surface with a radius of 76.5 mm, which we term *observation sphere* in the following and which was embedded in the middle of the GM layer. The triangles of the observation sphere had a mean edge length of 1.9 mm. FEM fields were evaluated by performing the FEM calculations and afterwards interpolating the electric fields on the *observation sphere* nodes using the SPR procedure.

**Figure 1:**
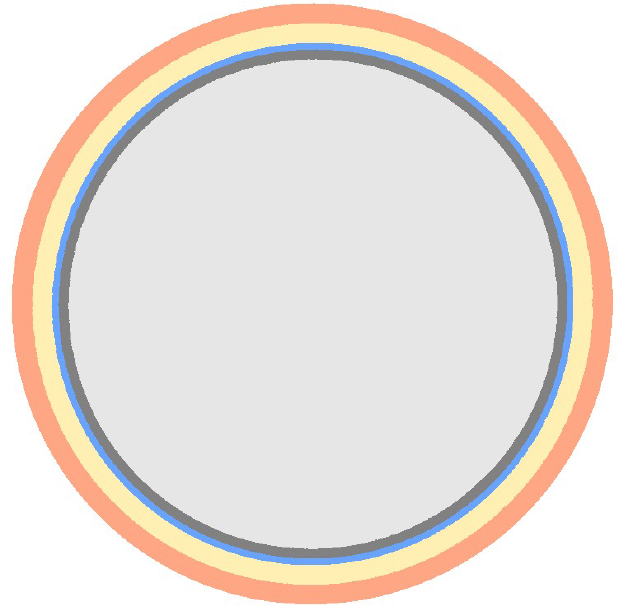
Phantom with five concentric spherical shells emulating head tissues.

**Table 1:** Number of Nodes, density of nodes in the spherical surfaces, and mean edge length for the tetrahedral elements in the spherical phantoms.

| Number of nodes ($\times 10^5$) | Number of tetrahedra ($\times 10^6$) | Mesh density, nodes/mm<sup>2</sup> | Mean edge length, mm |
| --- | --- | --- | --- |
| **0.46** | 0.26 | 0.07 | 5.1 |
| **1.24** | 0.72 | 0.14 | 3.6 |
| **3.03** | 1.80 | 0.28 | 2.7 |
| **8.40** | 5.05 | 0.55 | 1.9 |
| **26.97** | 16.43 | 1.21 | 1.3 |
| **72.69** | 44.63 | 2.41 | 0.92 |

We used the analytical solution developed by Sarvas [35] to validate our TMS FEM implementation. This solution gives the electric field produced by a magnetic dipole placed outside the sphere and is valid for any spherically symmetric conductivity distribution within the sphere. For the calculations, we positioned a magnetic dipole 10 mm above the surface of the sphere phantom and oriented it tangentially to the surface. In the FEM model, the conductivities of the two inner-most layers were set to 0.33 S/m, and the remaining conductivities were set to 1.79 S/m, 0.01 s/m and to 0.43 S/m (stated from inside to the outside [36]). Please note that the detailed choice of the conductivities is not important for the purpose of model validation, as long as they fall roughly in the range relevant for the later applications.

For the TES FEM implementation, we used the formula developed by Rush and Driscoll [37] for calculating the electric potentials caused by point electrodes in three concentric spherical shells. We used the same spherical models as for TMS but assigned to the three innermost shells a conductivity value of 0.33 S/m, making it into effectively a single shell. The model proposed by Rush and Driscoll features unrealistic point-wise electrodes. For this reason, instead of applying the electrodes directly to the surface of the spherical phantom, we choose to simulate the electric field in an “extended” spherical shell model with an outer radius of 102 mm. We used the extended phantom to calculate the electric potentials at the outer surface of the original spherical phantoms, where we used the results as Dirichlet boundary conditions for the FEM calculations. Our aim was to mimic the boundary conditions caused by small (EEG-sized rather than standard TES-sized) electrodes as a worst-case test, as the numerical accuracy of ensuring the set boundary conditions is more difficult to guarantee for small surface areas with a low number of nodes. Visual inspection of the potentials in the model’s outer layer confirmed that this procedure corresponds approximately to simulating a circular electrode of 12 mm diameter. As the equations proposed by Rush and Driscoll give the electric potential, we calculated the electric field by numerically evaluating potential differences between each of the sampling positions and positions 10^−4^ mm away along each direction.

Using the results obtained with the analytical models as reference, we assessed the error of the FEM solution in the *observation sphere* as

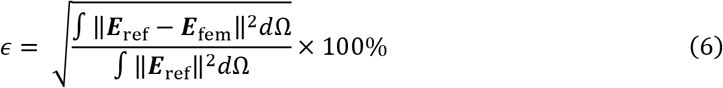

where ***E***_ref_ is the reference electric field on the observation sphere, and ***E***_fem_ the electric field obtained from the FEM calculations in SimNIBS.

### 2.4 Comparison with SimNIBS 2.1

We compared the new FEM implementation with SimNIBS 2.1, which is a popular software package for calculating TES and TMS electric fields [11] and which uses GetDP [38] to form and solve the FEM system. GetDP is a general environment for FEM problems, offering great flexibility in setting up and solving a large range of PDEs. In SimNIBS 2.1, GetDP is configured to use the CG method with a relative error of 10^−10^ and the incomplete cholesky preconditioner with two factor levels. Our aim was to confirm that the new implementation produces the same results as SimNIBS 2.1, and to quantify the improvements in computational efficiency and memory consumption.

We used a realistic head model created from automatically segmented magnetic resonance (MR) T1 and T2-weighted anatomical images (see [6] for details regarding the image acquisition parameters). The model was created in SimNIBS 2.1 using the *headreco pipeline* [39]. It has six tissue types corresponding to WM, GM, CSF, skull, skin and eyes (Figure 2a), and accounts for the major air cavities by sparing them in the mesh (effectively treating them as non-conducting vacuum). Tissue conductivities were set to 0.126 (WM), 0.275 (GM), 1.654 (CSF), 0.01 (skull), 0.465 (scalp) and 0.5 (skin) S/m, respectively [6]. The model has 6.8 × 10^5^ nodes, 3.5 × 10^6^ tetrahedra, average edge length of 2.15 mm, and a nominal average surface node density of 0.5 Nodes/mm^2^, which is the default setting for *headreco*.

**Figure 2:**
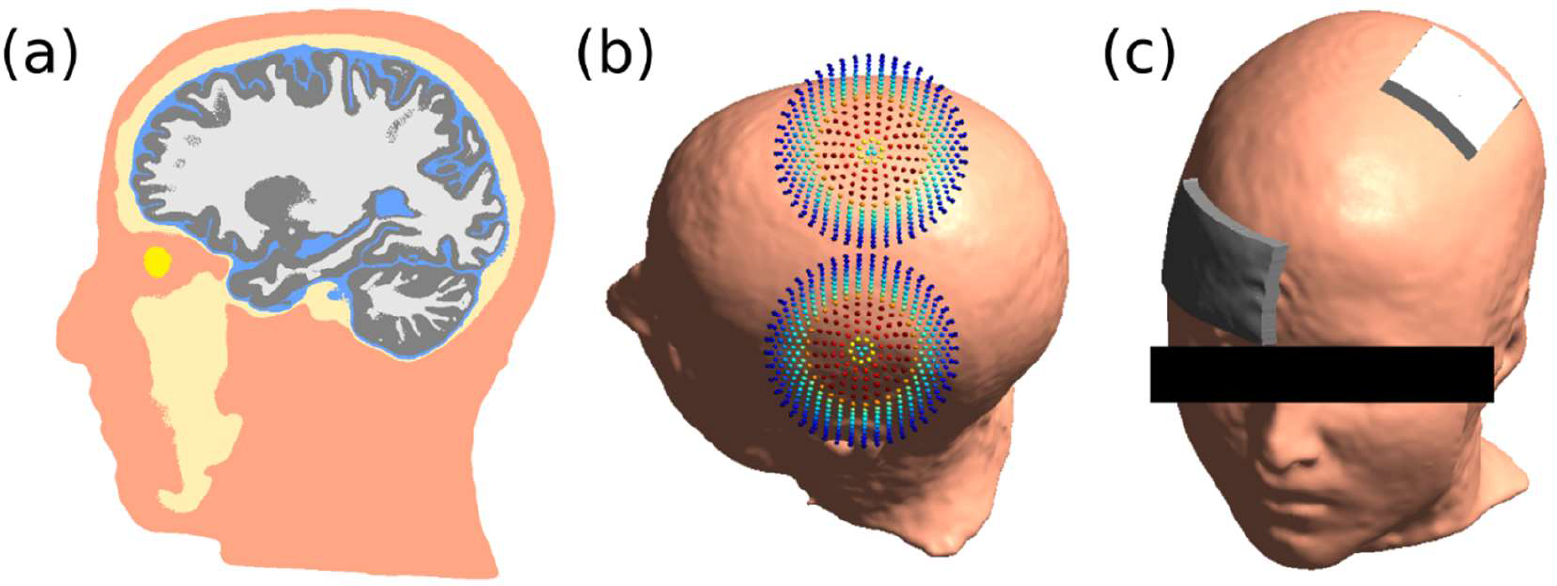
(a) Head Model with six tissues: white matter (WM, white), grey matter (GM, gray), cerebrospinal fluid (CSF, blue), skull (beige), skin (pink) and eyes (yellow). (b) Coil in TMS simulations. (c) Electrode montage for TES simulations.

In the TMS simulation, we calculated the electric field induced by a MagStim 70mm coil over the motor cortex, as shown in Figure 2(b). The current flux (*dI*/*dt*) was set to 1 *A*/*μs*. In the TES simulations, we calculated the electric field caused by a 50×50mm anode placed over the motor cortex and a 70×50mm supraorbital cathode, as shown in Figure 2(c). The current flux was set to 1 mA, and the electrode conductivity was set to 1 S/m. The potential in the electrodes was set homogeneously in the entire electrode surface, which corresponds to having a highly conductive material (such as a metal mesh) on the upper layer of the electrode.

### 2.5 Impact of Head Mesh Discretization on Simulation Accuracy

We evaluated the convergence with increasing mesh density for a realistic head mesh in order to provide estimates for the error caused by the discretization process. SimNIBS 2.1 uses the *headreco* pipeline to segment the MRI volumes into tissue classes, reconstruct surfaces that represent the tissue boundaries from the segmentations and create a volumetric tetrahedral mesh from the surfaces. Mesh density is controlled by setting the node density of the reconstructed surfaces (measured as nodes/mm^2^), which are than resampled using *meshfix* [40]. Here, we systematically varied the surface node density to control the anatomical accuracy of the head model and therefore obtain an insight into the node density that is required to guarantee a sufficient accuracy of the simulation results.

We constructed five head models with varying node densities based on the same MR segmentations (Table 2). Figure 3 shows in the left column a part of the GM surface around the left motor cortex and on the right column a cut of the CSF, GM and WM volumes in the same region, for each head model. Increasing node density leads to smaller tetrahedral elements, but also to an anatomically more accurate representation of the sulci. Both effects contribute to increase the accuracy of the simulated fields. In order to decouple them, we have in addition refined the four least dense head meshes by splitting their constituent tetrahedral elements. These head meshes have the same anatomical accuracy in representing the tissues, but the much finer elements allow for more accurate FE basis functions. The properties of these meshes are listed in Table 3.

**Figure 3:**
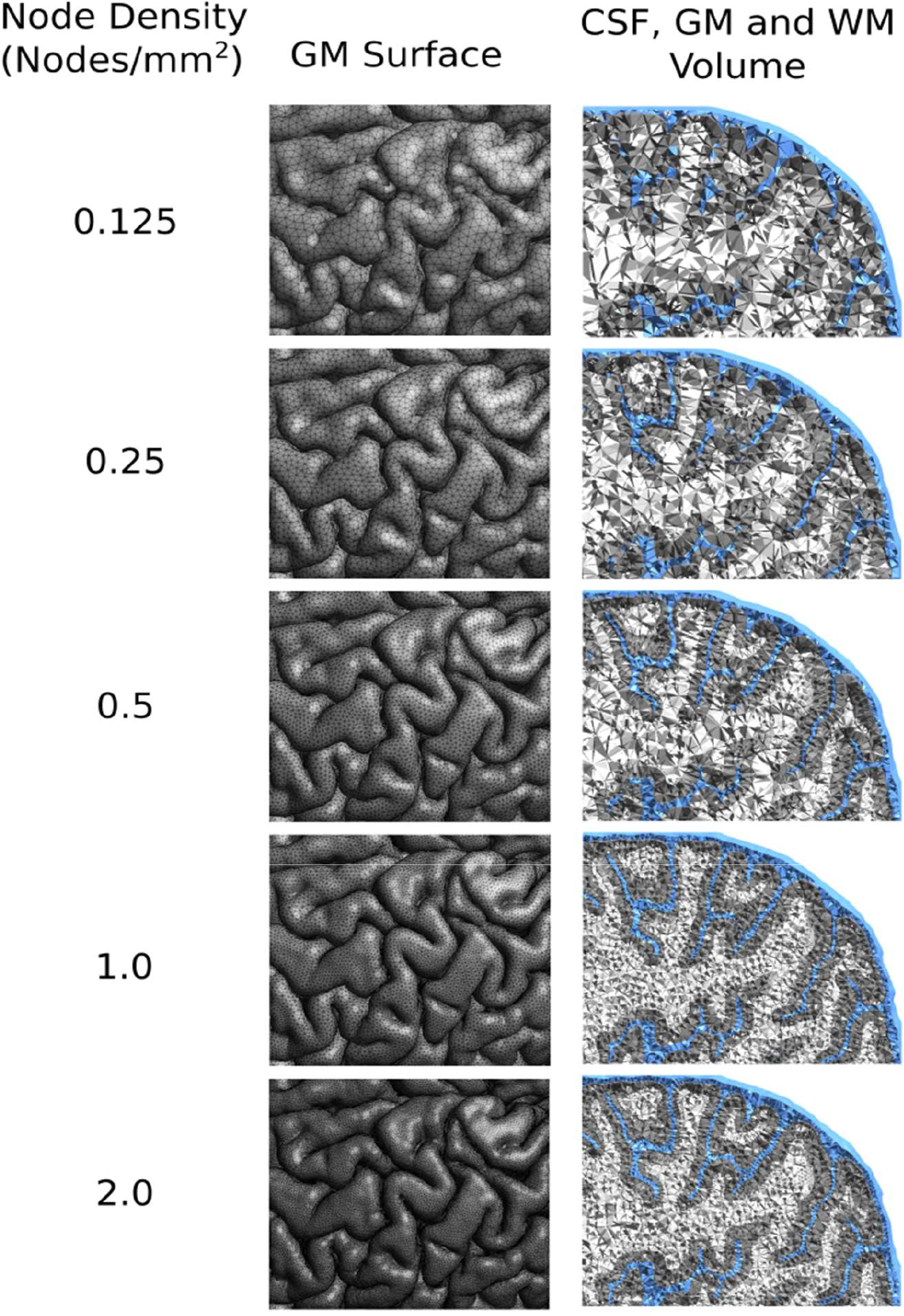
GM surface and GM, WM and CSF volumes at increasing mesh resolutions. Notice that not only the elements become finer, but also that the anatomical details of the CSF/GM boundary in the sulci are better preserved at higher mesh resolutions.

**Table 2:** Nominal and achieved node densities in the surfaces, number of nodes, number of tetrahedra and mean edge length in the tetrahedral elements for the five head models of different resolutions.

| Nominal Density<br>(Nodes/mm<sup>2</sup>) | Achieved Density<br>(Nodes/mm<sup>2</sup>) | Number of<br>nodes ($\times 10^5$) | Number of<br>tetrahedra<br>($\times 10^6$) | Mean edge<br>length, mm |
| --- | --- | --- | --- | --- |
| **0.125** | 0.149 | 1.3 | 0.7 | 3.63 |
| **0.25** | 0.275 | 3.0 | 1.7 | 2.28 |
| **0.5** | 0.520 | 6.8 | 3.5 | 2.15 |
| **1.0** | 0.996 | 20 | 11.2 | 1.48 |
| **2.0** | 2.001 | 33 | 18.5 | 1.23 |

**Table 3:** Mesh density data for the refined versions of the head meshes

| Original Density<br>(Nodes/mm<sup>2</sup>) | Refined mesh<br>Density<br>(Nodes/mm<sup>2</sup>) | Number of<br>nodes in<br>refined<br>mesh($\times 10^5$) | Number of<br>tetrahedra in<br>refined mesh<br>($\times 10^6$) | Mean edge<br>length in<br>refined<br>mesh, mm |
| --- | --- | --- | --- | --- |
| **0.125** | 0.592 | 10.2 | 5.9 | 1.88 |
| **0.25** | 1.142 | 23.6 | 13.7 | 1.46 |
| **0.5** | 2.087 | 48.8 | 28.4 | 1.11 |
| **1.0** | 4.036 | 152.3 | 89.4 | 0.78 |

We used the same TMS and TES set-ups as described in Section 2.4 (Fig. 2) to compare the fields across the 5 mesh densities. For that, we interpolated the electric fields on the nodes of a surface located in the middle of the gray matter layer, obtained through CAT12 (http://www.neuro.uni-jena.de/cat) during the *headreco* segmentation. This surface has 2.5 × 10^5^ nodes, a node density of 1.2 nodes/mm^2^ and a mean edge length of 1.0 mm). Interpolations were performed using either the unprocessed, elementwise-constant electric fields or the fields recovered through the SPR procedure.

### 2.6 Sulcus Phantom

We calculated the electric field in a simplified representation of a sulcus (Figure 4a) in order to explore the influence of element size and anatomical features on the numerical accuracy in more detail. As this model has an analytical geometric description, this approach allowed us to keep a good anatomical fidelity of the surfaces while changing element size. The model was composed of bone, CSF, GM, and WM. The model dimensions were set to mimic those of a real sulcus, and then sufficiently extended along the X and Z dimensions to ensure that the vertical boundaries did not have a dominant influence on the field distribution in the center of the phantom. Conductivities were set to the same values as used the head models.

**Figure 4:**
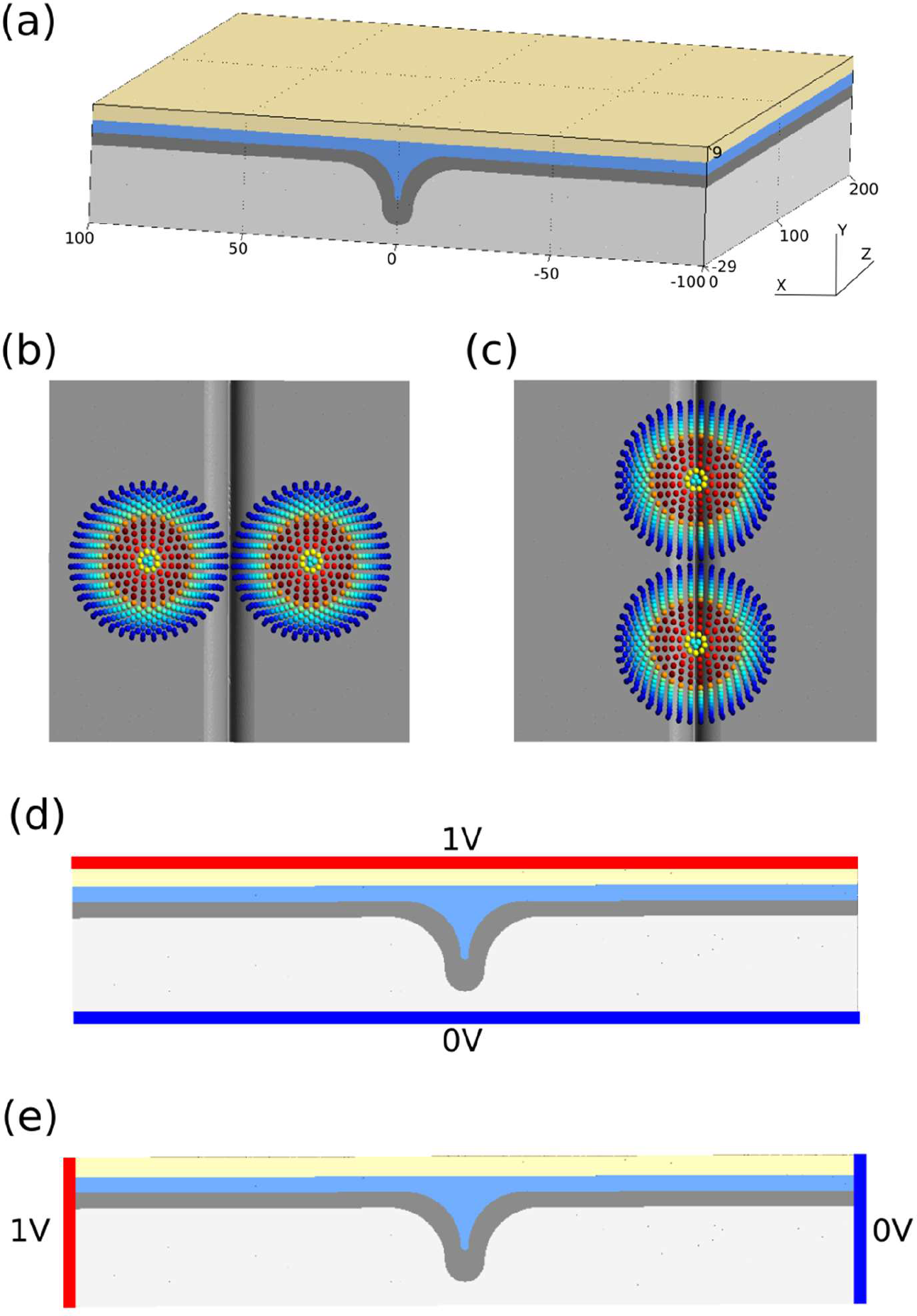
(a) Gyrus phantom with skull (beige), CSF (blue), GM (grey) and WM (white). Dimensions are shown in mm. (b) TMS simulation with the strongest fields induced parallel to the sulcus. (c) TMS simulation with the strongest fields induced perpendicular to the sulcus. (d) TES simulation with the potentials set at the horizontal boundaries at the top and bottom of the model. (e) TES simulation with the potentials set at two of the horizontal sides.

We simulated two positions for the TMS coil, with the coil being positioned 40 mm above the bone layer and oriented perpendicular (Figure 4b) or parallel (Figure 4c) to the sulcus. We also evaluated two cases in which we aimed to roughly mimic the field of an unfocal TES set-up. For that, we set potentials along the upper and lower boundaries of the model (Figure 4d) or along its left and right vertical boundaries (Figure 4e). For each case, we simulated the field in six models with increasing resolution (Table 4).

**Table 4:** Number of Nodes, density of nodes in the gray matter surface and mean edge length of the tetrahedral elements for the sulcus models.

| Number of nodes ($\times 10^5$) | Number of tetrahedra ($\times 10^6$) | Mesh density, nodes/mm<sup>2</sup> | Mean edge length, mm |
| --- | --- | --- | --- |
| **0.51** | 0.26 | 0.13 | 3.8 |
| **0.83** | 0.43 | 0.18 | 3.2 |
| **1.5** | 0.77 | 0.30 | 2.7 |
| **3.0** | 1.6 | 0.53 | 2.0 |
| **8.2** | 4.4 | 1.16 | 1.4 |
| **17.2** | 9.4 | 2.1 | 1.1 |

We visualized the field and numerical errors in a region of interest of the gray matter contained in an 80 mm x 20 mm box along the X and Z directions, placed in the center of the model. For each simulation, we interpolated the electric fields at the positions corresponding to the barycenters of the tetrahedra of the highest resolution model using the SPR procedure. Similarly as for the head models, we calculated the errors as the norm of the difference between the interpolated electric fields and the fields of the reference model.

## 3 Results

### 3.1 Validation in a Spherical Phantom

Figure 5 shows the error in the TMS electric field and TES electric potential and TES electric field for the six sphere models with increasing mesh density. The errors monotonically decay and are below 1% in the finest model investigated, validating the FEM implementation. The TES electric field has a larger error than the TMS field. We believe this is because the TES electric field is solely determined by the gradient of the potential, calculated using FEM, while the TMS electric field is also strongly dependent on the magnetic vector potential field which is calculated with a higher accuracy due to its simple analytical form.

**Figure 5:**
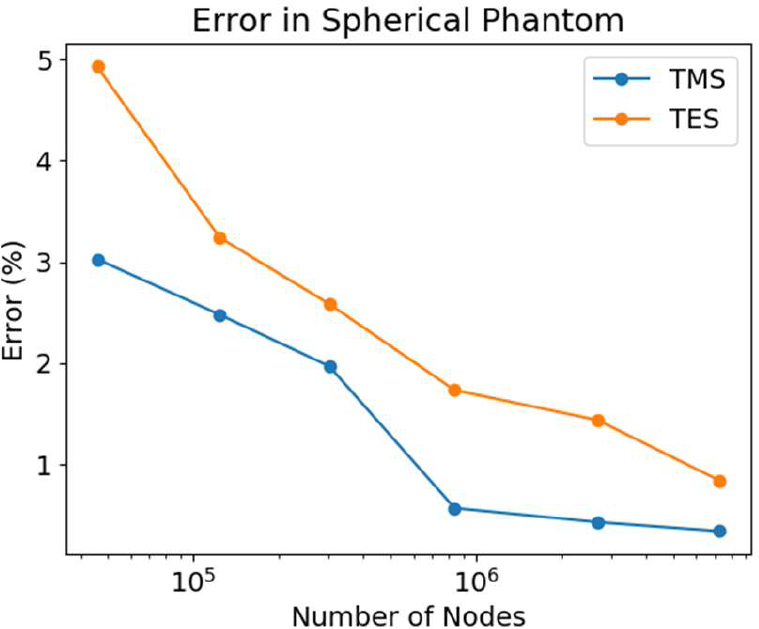
Error in the TMS and TES electric fields simulations for spherical phantoms with different number of nodes. The FEM results converge with increasing mesh density and show good agreement with the analytical solutions.

### 3.2 Comparison with SimNIBS 2.1

We used Equation 6 to compare the electric field in the GM volume obtained with SimNIBS 2.1 and the new FEM implementation and using the solution obtained with 2.1 as a reference. We saw relative differences of 0.052% and 0.046% for TMS and TES, respectively, indicating that SimNIBS 2.1 and 3.0 yield very similar results.

Figure 6 shows the time and the memory requirements assembling and solving a FEM system in SimNIBS 2.1 and the new implementation (SimNIBS 3.0). The calculations were performed on a laptop computer with an Intel i7-7500U processor (2 cores, 4 threads), 16 GB of memory, a SSD as hard-drive and running Ubuntu Linux 18.04. Times are given as the average ± standard deviation of ten runs. The new FEM approach is almost ten times faster for assembling and solving a TMS FEM system (222.0 ± 13.0 versus 23.0 ± 0.8 seconds) and almost five times faster for assembling and solving a TES FEM system (98.7 ± 4.6 versus 21.8 ± 1.2 seconds). Memory consumption is reduced by a factor of almost four.

**Figure 6:**
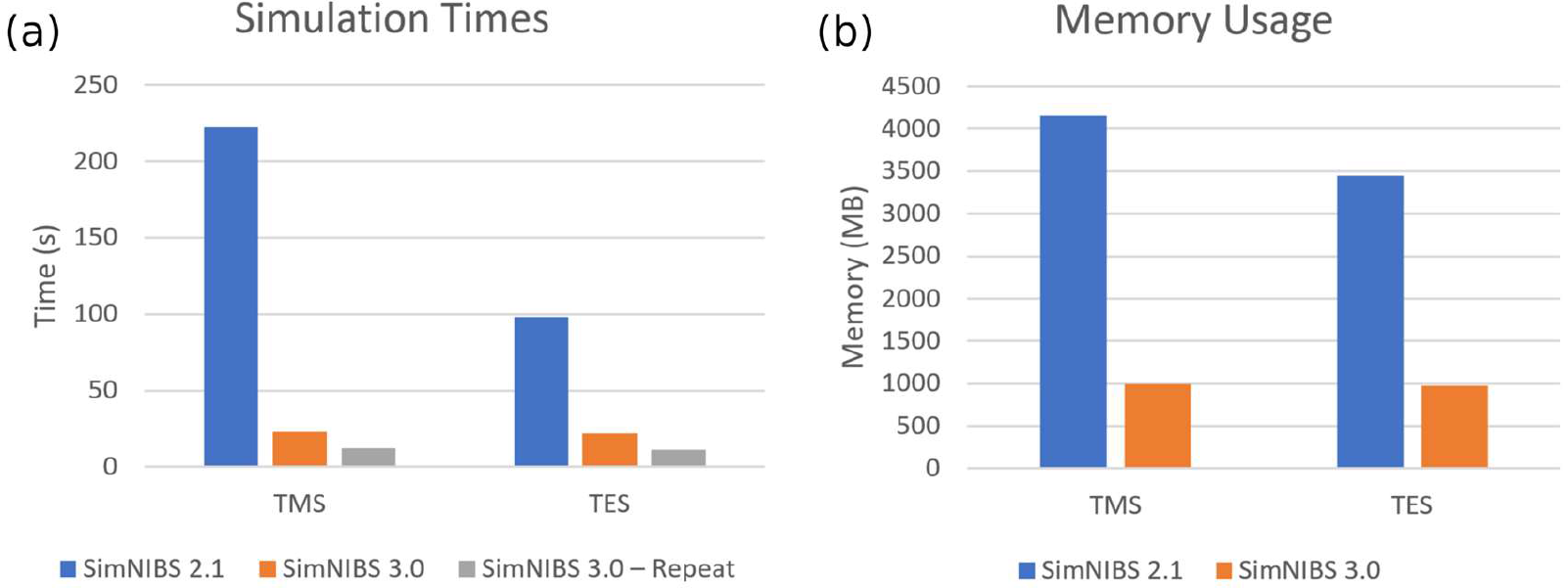
(a) Time to assemble and solve a FEM system and (b) memory usage in SimNIBS 2.1 and SimNIBS 3.0. SimNIBS 3.0 can reuse stiffness matrices and preconditioners, yielding a large speed up in specific applications where many simulations with the same mesh are needed.

In scenarios where the head mesh, tissue conductivities and the Dirichlet boundary conditions are kept constant, such as when running simulations with changing coil positions or construction of leadfield matrices for TES optimization [14], the new implementation re-utilizes the stiffness matrices and preconditioners, leading to further speed up. We observed that consecutive simulations take 11.7 ± 0.5 seconds for the TMS system and 11.0 ± 0.9 seconds for the TES system. This means that SimNIBS 3.0 can perform around 15 TMS field calculations in the same time that SimNIBS 2.1 uses to calculate a single simulation, while spending less of the system’s memory.

To perform a whole electric field simulation, SimNIBS goes through more steps than just the FEM, such as interpolating the *∂****A***/*∂t* field in TMS [41], meshing electrodes in TES, and calculating the gradient of the electrical potentials. When considering all those steps, the speed up is of around seven times for a single TMS simulation (235.31 ± 13.6 versus 33.0 ± 1.0) and around 2.5 times for a single TES simulation (162.2 ± 4.4 versus 60.2 ± 3.1 seconds). Notice that other parts of the code beside the FEM have also been optimized.

### 3.3 Impact of Head Mesh Discretization on Simulation Accuracy

Figure 7 shows the electric field on this middle cortical surface for the different mesh resolutions, interpolated using the SPR procedure. It also shows the errors, defined as the norm of the difference between the electric field obtained at each of the four lower resolutions and the field obtained at the highest resolution, at each position.

**Figure 7:**
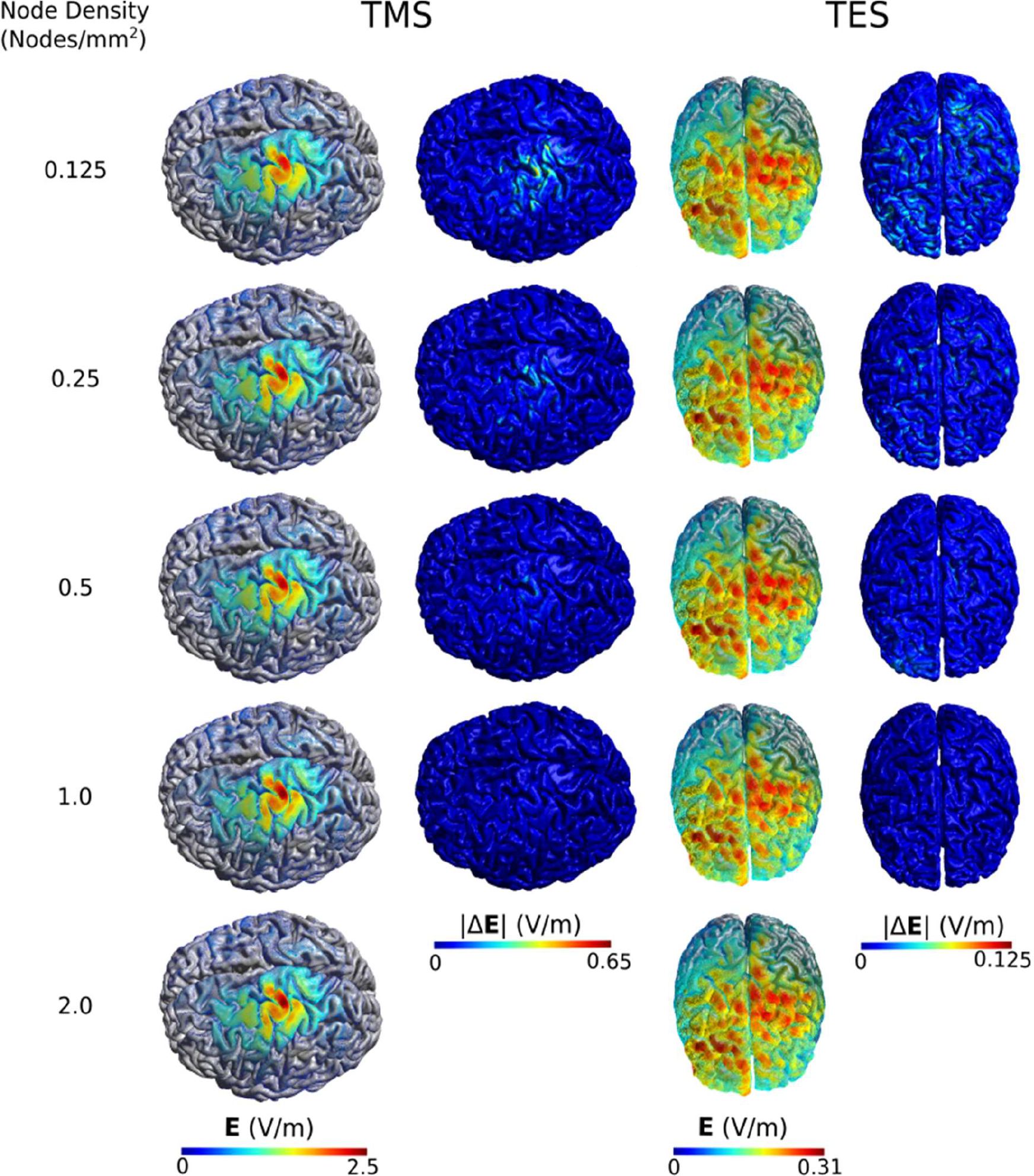
Electric field **E** and the error relative to the highest resolution model |Δ**E**| in the middle cortical layer for the TMS and TES simulations.

Even the results obtained for the lowest mesh density of 0.125 nodes/mm^2^ give a good overview of the overall field distribution in GM, suggesting that this density might be adequate for qualitative visualizations. However, the absolute errors in the estimated field strengths are still high (see below for further details). At 0.5 nodes/mm^2^, the values for the maxima of the electric fields seems to be better resolved, indicating this model is better suited for quantitative analysis.

We used Equation 6 to quantify the errors with the electric field obtained with the highest resolution model as ***E***_ref_ (Fig. 8a). In order to understand whether the improvement in simulation accuracy with increasing mesh resolution was primarily driven by the reduced size of the tetrahedral elements, or by the better anatomical fidelity in representing the GM/CSF and other tissue boundaries, we also evaluated the error on the models refined by splitting (described in Section 2.5).

**Figure 8:**
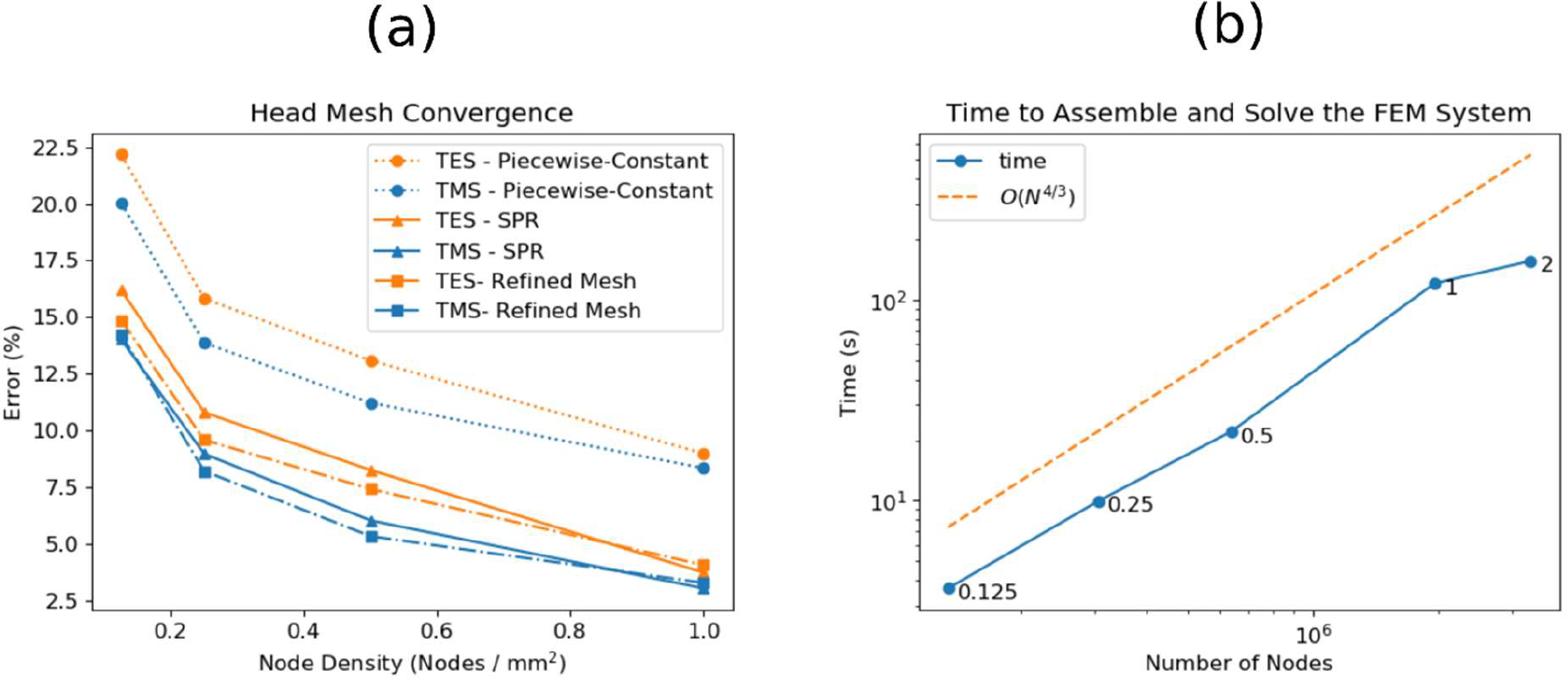
Convergence of the electric field in the middle cortical layer for TMS and TES simulations (a), and time to assemble and solve FEM systems (b). Errors were accessed using Equation 6, with the electric field obtained with 2 nodes/mm2 model as a reference.

In addition, in order to evaluate the effect of the interpolation method, we used both the SPR-based interpolation and the unprocessed, element-wise constant electric fields. The former was used in both the original and the refined mesh, and the later was only used in the original mesh. This comprehensive analysis allows us to identify the main factors underlying simulation accuracy. Finally, we also assessed how the time for the FEM calculations depended on the mesh resolution.

Generally, errors were high at the lowest mesh density, clearly exceeding 10% for all cases. Employing the SPR-based interpolation substantially improved the field estimates and helped to achieve errors around 6% for TMS and 8% for TES for a mesh density of 0.5 nodes/mm^2^. This is the standard mesh density in the *headreco* pipeline and was selected as tradeoff between acceptable simulation time and numerical error. Interestingly, refining the head mesh by splitting only marginally improved accuracy. In fact, the refined version of the mesh with 0.25 nodes/mm^2^ has more nodes than the head model with 1.0 nodes/mm^2^, but an error that is approximately two times larger. This indicates that the key factor for convergence is the anatomical fidelity of the modelled tissue surfaces, rather than only element size. In line with the results obtained for the spherical phantom, errors are lower for the TMS simulations compared to the TES simulations.

Direct pairwise comparisons of the original and refined versions of the head meshes (section S.4 of the supplementary material) reveals an expected clear improvement in numerical accuracy for the finer mesh. However, such differences are not present here when comparing to an anatomically more accurate head model as reference, i.e. a better numerical accuracy alone does not seem to translate into a better overall accuracy. This suggests again that anatomical fidelity and element size are both important factors that have to be commonly considered to obtain accurate simulations.

The times to assemble and solve the FEM system are shown in Figure 8B. As the TES and TMS simulation times are very similar, only the simulation times for TMS are shown as a function of the number of nodes in the mesh. The simulation times grow approximately with 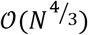 where *N* is the number of nodes in the mesh. This is what is expected from a CG solver for 3D elliptical problems [42].

### 3.4 Sulcus Phantom

The electric fields and the errors for the sulcal phantom are shown in Figure 9. Interestingly, the electric field distribution in the sulcus is highly dependent on the set-up, especially for TES. This is due to the large conductivity difference between CSF and GM (1.654 versus 0.275 S/m), which favors current flow through CSF. Thus, when the potentials are set along the top and bottom of the model, the current tend to flow to the bottom of the sulci, where they enter GM. On the other hand, when the potentials are set on the sides of the model, the currents will tend to flow along the CSF layer on the top of the phantom. Generally, the errors tend to be concentrated in the regions of highest curvature at the top or at the bottom of the gyrus, and there is a large difference in the magnitude of the errors across the different stimulation set-ups.

**Figure 2:**
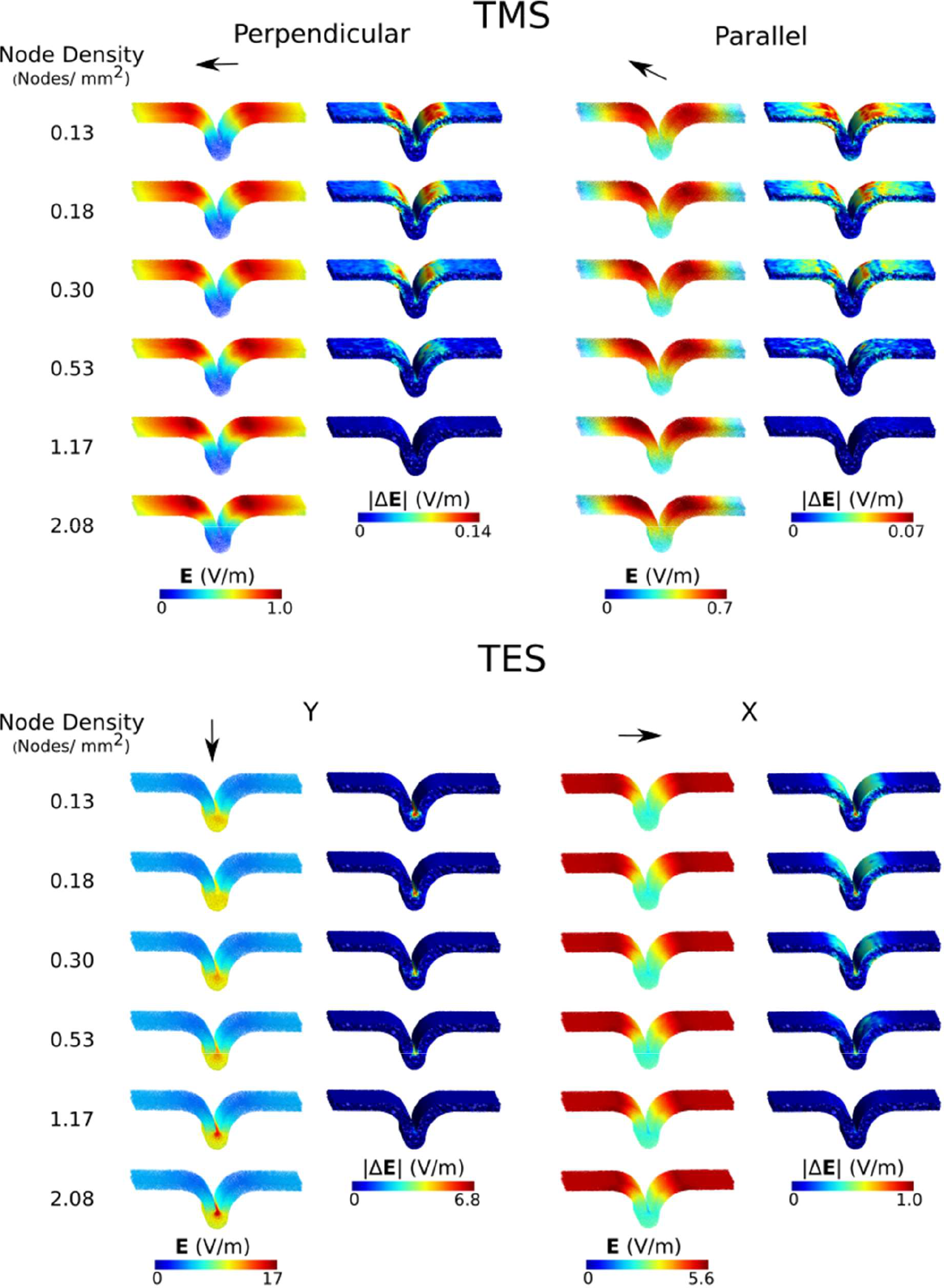
Electric fields and errors in the region of interest of the sulcus phantom. The arrows indicate the main direction of the electric field vectors, and the errors were calculated in relation to the highest resolution model.

Figure 10 shows the average numerical error in the region of interest, calculated using Equation 6 and in relation to the highest resolution model. As in the previous cases, the errors steadily decrease as the model in refined, indicating convergence. As expected from Figure 9, the errors for the TES simulations with the potentials set at the horizontal boundaries are much larger than for the other three set-ups. This is because the electric currents tend to flow along the conductive CSF layer, entering the grey matter at the bottom of the sulcus, where it spreads out again. This effect causes the electric field to be highest in the gray matter of the sulcus fundus and to vary strongly within this relatively small region, so that a higher mesh resolution is required to accurately capture this effect.

**Figure 3:**
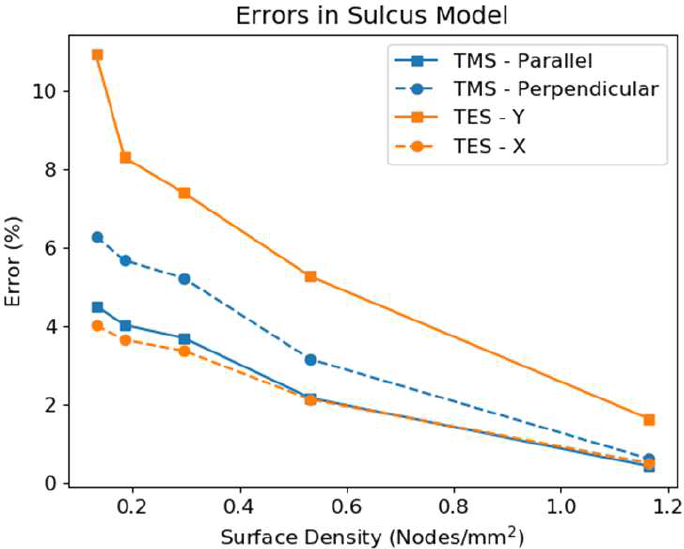
Errors in the electric field as a function of node density, in relation to the highest resolution model.

## 4. Discussion and Conclusion

In the previous sections we presented an efficient implementation for performing electrical field simulations based on FEM for both TMS and TES. This implementation will be available as a part of SimNIBS version 3.0. Validation of the new implementation show that electric fields obtained agree well with analytical solutions of the field in spherical conductors and are very similar to electric fields obtained with a previous implementation (SimNIBS 2.1). However, the new implementation clearly outperforms the previous implementation, runs simulations up to nine times faster and requires up to four times less memory.

Our validations show that TMS and TES simulations converge with increasing anatomical fidelity of the GM layer. This suggests that a detailed representation of the cortical folding is important to obtain accurate results. Furthermore, our results indicate that estimating the elementwise-constant electric field obtained directly from first-order FEM may require a very high node density in order to ensure adequate accuracy, but that this can be somewhat compensated by the use of SPR interpolation. It is also interesting to notice that, while refining the head mesh does increase the numerical accuracy of the solution (e.g. Supplementary Fig. S.1), this does not necessarily improve the overall accuracy when compared to a reference model that is also anatomically more accurate. This shows that increasing the overall model accuracy requires improving both the anatomical fidelity and the density of the head mesh.

The data suggests that the choice of an appropriate value for the mesh density is highly dependent on the application. Values as low as 0.25 nodes/mm^2^ might be appropriate for qualitative analysis of the electric field distributions. A density of 0.5 nodes/mm^2^ is currently set as standard in SimNIBS and seems to give an acceptable trade-off between efficiency and accuracy, with overall errors in the range of 6% to 8% and numerical errors below 5%. Values of 1.0 nodes/mm^2^ or higher are needed to ensure overall errors being consistently lower than 5%, as e.g. required for careful analyses of the field in the sulci, which are not properly captured in low resolution models. The increased computational efficiency of the new FEM implementation ensures that simulations with these high node densities are no longer prohibited by the increased computational efforts, and can now easily be used on a regular basis. As the errors are consistent across stimulation modalities, we do not expect them to exhibit very large variations across simulation set-ups, or between head models of healthy individuals, given that the segmentation is accurate.

We believe that overall simulation errors in the observed range are currently still acceptable, given that other error sources such as segmentation errors [39], the uncertainty of the values of the ohmic tissue conductivities [43], putative inaccurate or simplified modeling of the of TMS coil or TES electrode properties and positions [7] can cause errors in or above this range. For example, in a recent study [43], we found the uncertainty in peak electric values due to uncertainty in tissue conductivities to be around 5% of the mean value in TMS, but around 20% of the mean value in TES. This suggest that future efforts to increase simulation accuracy will have to commonly account for all these factors, rather than focusing on improving the FEM accuracy alone. In addition, considering the state of knowledge on how the electric field modulates neural activity, simulation errors in the reported range do currently not affect the conclusions that can be drawn from the simulation results.

In a related manner, this study is limited to FEM with first order tetrahedral elements. Changing to higher order elements, or employing alternative methods to CG-FEM (as used in the current work), such as DG-FEM [36,44], BEM [45,46], and BEM-FMM [34,47] will likely help to obtain faster convergence in simple geometries such as the sphere and the sulcus phantoms. However, for all these methods, the results can still only be accurate as far as the head model is accurate, meaning that we still expect error bounds similar to those displayed in Figure 8 to hold when compared to a reference model with better anatomical fidelity of the GM sheet. In line with our results, Engwer et. al. [36] noted in the context of DG-FEM for the EEG forward problem that the errors in the solutions seems to be dominated by errors in adequately representing the anatomy, rather than by the solution method (DG-FEM vs. CG-FEM).

In combination, the results for the full head model and the sulcus model indicate that the numerical accuracy of the solutions benefits from an accurate anatomical representation of the tissue boundaries. In particular, the sulcus model shows that a higher mesh density around strongly curved parts of the GM/CSF boundary, where the electric potential can show relative abrupt changes, is helpful in order to avoid local simulation errors around those parts. One course of action suggested by our results would be to improve the surface meshing such that it can preserve the sulci better and increase the node density in regions of high curvature. This could be done by changing the resampling algorithm such that it takes into account local curvature or by other methods such as harmonic maps [48], or isoperimetric higher order elements. Those methods would hopefully allow us to obtain accurate representations of the tissue surface while keeping a small number of elements, and thus improve simulation accuracy at a low computational cost.

In summary, we present a new FEM implementation, which will be made available in SimNIBS 3.0. This implementation leads to up to twenty times faster simulations than in SimNIBS 2.1. This will provide a massive speed up in applications requiring repeated simulations such as calculating several different TMS positions, obtaining leadfields for TES optimization, or performing uncertainty quantification. We also estimated the accuracy for the electric field simulations, demonstrating that that anatomical fidelity is at least of similar importance as mesh density in order to obtain accurate results.

## Acknowledgements

This work was suppoted by Lundbeckfonden (grant Nr. R118-A11308), and NovoNordisk fonden (grant Nr. NNF14OC0011413).

