## Supplementary Material for "Efficient Electric Field Simulations for Transcranial Brain Stimulation"

[1]Danish Research Centre for Magnetic Resonance, Centre for Functional and Diagnostic Imaging and Research, Copenhagen University Hospital Hvidovre, Denmark
[2]Department of Health Technology, Technical University of Denmark, Kgs. Lyngby, Denmark
[3]Department of Applied Mathematics and Computer Science, Technical University of Denmark, Kgs. Lyngby, Denmark
[*]Equal Contribution

February 4, 2019

### S.1 Weak Form of the Differential Equations

#### S.1.1 TES

In TES, a Laplace equation

$$\begin{aligned}
\nabla \cdot (\underline{\sigma} \nabla \phi) &= 0 \quad \text{in } \Omega, && \text{(S.1)}\\
\boldsymbol{J} &= -\underline{\sigma} \nabla \phi \quad \text{in } \Omega, && \text{(S.2)}\\
\phi &= 0 \quad \text{on } \Gamma_1, && \text{(S.3)}\\
\phi &= a \quad \text{on } \Gamma_2, && \text{(S.4)}\\
\boldsymbol{J} \cdot \hat{\boldsymbol{n}} &= 0 \quad \text{on } \partial\Omega \setminus (\Gamma_1 \cup \Gamma_2)\,, && \text{(S.5)}
\end{aligned}$$

governs the electric potentials $\phi$ in a region $\Omega$ with conductivity values $\underline{\sigma}$, represented by a 3x3 symmetric positive definite (SPD) tensor, which can vary spatially. The current density $\boldsymbol{J}$ can be calculated from the gradient of the electric potential and the conductivity values, and $\hat{\boldsymbol{n}}$ is a vector normal to the surface. In this setting, we assume that the current is injected through two surface electrodes, $\Gamma_1$ and $\Gamma_2$, but we can easily extend the equations to add more electrodes.

Conductivity values may suffer discrete jumps across tissue interfaces. Therefore, we also use the continuity conditions [1]

$$\phi' = \phi'', \tag{S.6}$$

$$\boldsymbol{J}' \cdot \hat{\boldsymbol{n}} = \boldsymbol{J}'' \cdot \hat{\boldsymbol{n}}, \tag{S.7}$$

where $\phi'$ is the electric potential and $\boldsymbol{J}'$ is the current density at one side of the interface and $\phi''$ and $\boldsymbol{J}''$ are the same quantities at the other side of the tissue interface.

To obtain the weak form of the partial differential equation (PDE), we introduce the test function $\nu \in V$, $V = \{\nu \in H^1(\Omega);\ \nu = 0 \text{ on } (\Gamma_1 \cup \Gamma_2)\}$, multiply it to Equation S.1 and integrate it over the domain $\Omega$ [2], obtaining

$$\int_\Omega \nabla \cdot (\underline{\sigma} \nabla \phi)\, \nu \; d\boldsymbol{x} = 0 \quad \forall \nu \in V, \tag{S.8}$$

where $\boldsymbol{x}$ represents a position in the domain $\Omega$. Using Gauss' theorem and the continuity conditions (Equations S.6 and S.7), we can re-write the equation above as

$$-\int_\Omega (\underline{\sigma} \nabla \phi) \cdot \nabla \nu \; d\boldsymbol{x} + \int_{\partial\Omega} (\underline{\sigma} \nabla \phi)\, \nu \cdot \hat{\boldsymbol{n}} \; dS = 0 \quad \forall \nu \in V. \tag{S.9}$$

Using the Neumann boundary condition (Equation S.5) as well as the definition of the space $V$, we can eliminate the surface integral and obtain

$$\int_\Omega (\underline{\sigma} \nabla \phi) \cdot \nabla \nu \; d\boldsymbol{x} = 0 \quad \forall \nu \in V. \tag{S.10}$$

Solutions to the weak form (Equation S.10) that fulfill the Dirichlet boundary conditions and are sufficiently regular also solve the strong form (Equation S.1) [2].

Using the Dirichlet boundary conditions, we assume that the potentials the electrodes are known. However, in TES we typically set the electric current through the electrodes, and not the electric potential. Therefore, after calculating the electric potential, we calculate the total current flow through the electrodes and use the linearity of the Laplace equation to re-scale the results according to the desired current. Another approach is to directly apply Neumann boundary conditions in the electrode $\Gamma_2$. This way, we have an approximation of the actual boundary conditions (which are of the Dirichlet type), but can skip the re-scaling step.

This is done by replacing the boundary condition in Equation S.4 by

$$\boldsymbol{J} \cdot \hat{\boldsymbol{n}} = \frac{I}{A(\Gamma_2)} \quad \text{on } \Gamma_2, \tag{S.11}$$

where $I$ is the electrical current and $A(\Gamma_2)$ the area of the electrode $\Gamma_2$. In this case, we obtain the weak form of the PDE

$$\int_\Omega (\underline{\sigma}\nabla\phi) \cdot \nabla\nu \; d\boldsymbol{x} = -\frac{I}{A} \int_{\Gamma_2} \nu \; dS \quad \forall \nu \in V. \tag{S.12}$$

#### S.1.2 TMS

For TMS, a Poisson equation governs the electric potential [3]

$$\nabla \cdot (\underline{\sigma}\nabla\phi) = -\nabla \cdot \left(\underline{\sigma}\frac{\partial \boldsymbol{A}}{\partial t}\right) \quad \text{in } \Omega, \tag{S.13}$$

$$\boldsymbol{J} = -\underline{\sigma}\nabla\phi - \underline{\sigma}\frac{\partial \boldsymbol{A}}{\partial t} \quad \text{in } \Omega, \tag{S.14}$$

$$\boldsymbol{J} \cdot \hat{\boldsymbol{n}} = 0 \quad \text{on } \partial\Omega, \tag{S.15}$$

where $\boldsymbol{A}$ is the magnetic vector potential created by the coil. Again, we have the continuity conditions given by Equations S.6 and S.7 [3]. To obtain the weak form, we introduce a test function $\nu \in V$, $V = H^1(\Omega)$, multiply it with Equation S.13 and integrate it over the domain $\Omega$

$$\int_\Omega \nabla \cdot \left(\underline{\sigma}\nabla\phi + \underline{\sigma}\frac{\partial \boldsymbol{A}}{\partial t}\right) \nu \; d\boldsymbol{x} = 0 \quad \forall \nu \in V. \tag{S.16}$$

We apply Gauss' theorem and the continuity conditions, obtaining

$$-\int_\Omega \left(\underline{\sigma}\nabla\phi + \underline{\sigma}\frac{\partial \boldsymbol{A}}{\partial t}\right) \cdot \nabla\nu \; d\boldsymbol{x} + \int_{\partial\Omega} \left(\underline{\sigma}\nabla\phi + \underline{\sigma}\frac{\partial \boldsymbol{A}}{\partial t}\right) \nu \cdot \hat{\boldsymbol{n}} \; dS = 0 \quad \forall \nu \in V. \tag{S.17}$$

Using the Neumann boundary condition (Equation S.15), we eliminate the surface integral, obtaining the weak form

$$\int_\Omega (\underline{\sigma}\nabla\phi) \cdot \nabla\nu \; d\boldsymbol{x} = -\int_\Omega \left(\underline{\sigma}\frac{\partial \boldsymbol{A}}{\partial t}\right) \cdot \nabla\nu \; d\boldsymbol{x} \quad \forall \nu \in V. \tag{S.18}$$

### S.2 Galerkin Method

Consider the weak forms of the PDEs (Equations S.10, S.12, S.18), which can be written in as

$$a(\phi, \nu) = l(\nu) \quad \forall \nu \in V. \tag{S.19}$$

For all three cases, we have

$$a(\phi, \nu) = \int_\Omega (\underline{\sigma}\nabla\phi) \cdot \nabla\nu \; d\boldsymbol{x}. \tag{S.20}$$

In the TES case with Dirichlet boundary conditions in both electrodes, we have

$$l(\nu) = 0. \tag{S.21}$$

In the TES case with Neumann boundary conditions in the electrode $\Gamma_2$, we have

$$l(\nu) = -\frac{I}{A(\Gamma_2)} \int_{\Gamma_2} \nu \, dS, \tag{S.22}$$

and, in the TMS case we have

$$l(\nu) = -\int_{\Omega} \left( \underline{\sigma} \frac{\partial \boldsymbol{A}}{\partial t} \right) \cdot \nabla \nu \, d\boldsymbol{x}. \tag{S.23}$$

Up until now, the problems are formulated in an infinite-dimensional functional space $V$. In order to solve the problem numerically, we need to define basis functions $\psi_i(\boldsymbol{x}),\ i = 1, \ldots, N$. The basis functions span a space

$$V^h = \bigcup_{i=1}^{N} V^i, \tag{S.24}$$

where $V^i$ is the space spanned by each basis function $\psi_i$. We will now look for solutions for PDEs in the finite-dimensional space $V^h$ by enforcing [2]

$$a^h(\phi^h, \nu^h) = l(\nu^h) \quad \forall \nu^h \in V^h. \tag{S.25}$$

We write the electric potential as a sum of basis functions

$$\phi^h(\boldsymbol{x}) = \sum_{i=1}^{N} \phi_i \psi_i(\boldsymbol{x}), \tag{S.26}$$

and substitute it in the bilinear form (Equation S.20), obtaining

$$a^h(\phi^h, \nu^h) = \sum_{i=1}^{N} \phi_i \int_{\Omega} (\underline{\sigma} \nabla \psi_i(\boldsymbol{x})) \cdot \nabla \nu^h \, d\boldsymbol{x}. \tag{S.27}$$

Substituting $\nu^h(\boldsymbol{x}) = \psi_j(\boldsymbol{x}),\ \forall\ j = 1, \ldots, N$, we obtain a system of $N$ equations that can be written as

$$\boldsymbol{S}\boldsymbol{u} = \boldsymbol{b}, \tag{S.28}$$

where $\boldsymbol{S}$ is the **stiffness matrix**. The entries of the matrix and vectors of the system are given by

$$S_{ij} = a^h(\psi_i, \psi_j), \tag{S.29}$$

$$u_i = \phi_i, \tag{S.30}$$

$$b_i = l^h(\psi_i). \tag{S.31}$$

It is easy to see that, in our case, the stiffness matrix is **symmetric**.

### S.3 The Finite Element Method

#### S.3.1 Basis Functions

For the FEM, we discretize the space $\Omega$ into simple non-overlapping geometric shapes called **elements**, thereby obtaining a space $\Omega^h$. The vertices of the elements are shared between many neighbors, and are called **nodes**. The basis functions $\psi_i(\boldsymbol{x})$ are typically piecewise-polynomial functions, defined at the nodes. In the 3D case, the most simple type of element is the linear tetrahedron. In this case, the basis functions are defined such that $\psi_i(\boldsymbol{x}_i) = 1$ in the position $\boldsymbol{x}_i$ of a given node $i$, and decays linearly within all elements containing the node $i$, reaching $\psi_i(\boldsymbol{x}_j) = 0$ at all nodes $j$ neighboring $i$. Within all elements that do not contain the node $i$, the basis function $\psi_i(\boldsymbol{x})$ is zero [4].

Using the basis functions, we can interpolate values of an arbitrary function $f(\boldsymbol{x})$ within a given element $\Omega^e$, given its values at the nodes of the element $\Omega^e$

$$f^e(\boldsymbol{x}) = \sum_{i\in\Omega^e} f_i\psi_i(\boldsymbol{x}) \quad \forall\boldsymbol{x} \in \Omega^e. \tag{S.32}$$

As the functions $\psi_i(\boldsymbol{x})$ are linear by definition, $f^e(\boldsymbol{x})$ is also linear and has the form

$$f^e(\boldsymbol{x}) = \begin{bmatrix} 1 & \boldsymbol{x}^T \end{bmatrix} \boldsymbol{\alpha} \quad \forall\boldsymbol{x} \in \Omega^e. \tag{S.33}$$

The coefficients $\boldsymbol{\alpha}$ are given by a $4 \times 1$ vector and can be determined by solving the system [4]

$$\begin{bmatrix} 1 & (\boldsymbol{x}_1^e)^T \\ 1 & (\boldsymbol{x}_2^e)^T \\ 1 & (\boldsymbol{x}_3^e)^T \\ 1 & (\boldsymbol{x}_4^e)^T \end{bmatrix} \boldsymbol{\alpha} = \boldsymbol{f}^e, \tag{S.34}$$

where $\boldsymbol{f}^e$ is a $4 \times 1$ vector with the values of the function in each of the tetrahedron nodes, and $(\boldsymbol{x}_i^e)^T,\ i = 1, 2, 3, 4$ are the element node positions.

#### S.3.2 Gradient Operator

From Equation S.33, we see that the local gradient of the function $f(\boldsymbol{x})$ in the element $\Omega^e$ is approximated by the three last indices of $\boldsymbol{\alpha}$

$$\nabla f^e(\boldsymbol{x}) = \begin{bmatrix} \alpha_2 \\ \alpha_3 \\ \alpha_4 \end{bmatrix}. \tag{S.35}$$

Those can be directly calculated by subtracting the first row of the system in Equation S.34, obtaining

$$\begin{bmatrix} f_2 - f_1 \\ f_3 - f_1 \\ f_4 - f_1 \end{bmatrix} = \begin{bmatrix} (\boldsymbol{x}_2^e - \boldsymbol{x}_1^e)^T \\ (\boldsymbol{x}_3^e - \boldsymbol{x}_1^e)^T \\ (\boldsymbol{x}_4^e - \boldsymbol{x}_1^e)^T \end{bmatrix} \begin{bmatrix} \alpha_2 \\ \alpha_3 \\ \alpha_4 \end{bmatrix}. \tag{S.36}$$

We can therefore define a local gradient operator $\boldsymbol{G}^e$

$$\nabla f^e(\boldsymbol{x}) = \boldsymbol{G}^e \boldsymbol{f}^e, \tag{S.37}$$

which is given by

$$\boldsymbol{G}^e = \begin{bmatrix} (\boldsymbol{x}_2^e - \boldsymbol{x}_1^e)^T \\ (\boldsymbol{x}_3^e - \boldsymbol{x}_1^e)^T \\ (\boldsymbol{x}_4^e - \boldsymbol{x}_1^e)^T \end{bmatrix}^{-1} \begin{bmatrix} -1 & 1 & 0 & 0 \\ -1 & 0 & 1 & 0 \\ -1 & 0 & 0 & 1 \end{bmatrix}. \tag{S.38}$$

#### S.3.3 Stiffness Matrix Assembly

Now, we can use the gradient operator to calculate entries of the stiffness matrix. From Equations S.27 and S.29, we have

$$S_{ij} = \int_{\Omega^h} (\underline{\sigma} \nabla \psi_i(\boldsymbol{x})) \cdot \nabla \psi_j(\boldsymbol{x})\ d\boldsymbol{x}. \tag{S.39}$$

We subdivide the integral over the domain $\Omega^h$ by integrals within each element $\Omega^e, e = 1, \ldots, N_e$, obtaining

$$S_{ij} = \sum_{e=1}^{N_e} \int_{\Omega^e} (\underline{\sigma} \boldsymbol{G}^e \boldsymbol{\psi}_i) \cdot \boldsymbol{G}^e \boldsymbol{\psi}_j\ d\boldsymbol{x}. \tag{S.40}$$

Where $\boldsymbol{G}^e$ is the gradient operator defined in Equation S.38. As we defined the basis functions $\psi_i(\boldsymbol{x})$ to be zero in elements $e$ that do not contain the node $i$, we have that $S_{ij} = 0$ if the nodes $i$ and $j$ are not neighbors. And, because each node has only few neighbors, the stiffness matrix is **sparse**.

To efficiently calculate the entries of the stiffness matrix, we introduce two mappings

- $k_i^e$ is a mapping from global node index $i$ to the local node index (numbered from 1 to 4) of the element $e$.
- $i_k^e$ is a mapping from local node index $k$ in the element $e$ to the corresponding global node index.

Because the basis functions $\psi_i$ have a value of 1 in their corresponding node and of 0 in all other nodes, the operation $\boldsymbol{G}^e \boldsymbol{\psi}_i$ corresponds to the $k_i^e$ column of the local gradient operator, $\boldsymbol{G}^e_{k_i^e}$. We therefore can calculate the entries of the stiffness matrix as

$$S_{ij} = \sum_{e | i,j \in \Omega^e} \int_{\Omega^e} \left( \underline{\sigma} \boldsymbol{G}^e_{k_i^e} \right) \cdot \boldsymbol{G}^e_{k_j^e}\ d\boldsymbol{x}. \tag{S.41}$$

Assuming that $\underline{\sigma}$ is constant or varies linearly within each element, we can calculate the integral in Equation S.41 exactly using only one integration point, in the middle of the element. In the current work, we always use piecewise-constant values for $\underline{\sigma}$, with the conductivity jumps always occuring between elements.

#### S.3.4 Assembly of the Right-Hand Side

##### S.3.4.1 TES with Neumann boundary conditions

Equation S.22 gives us the right-hand side (RHS) of a TES system with Neumann type boundary conditions. To calculate it, we need to define surface elements $\partial\Omega^e$, which in the case of tetrahedral volume elements are the triangles corresponding to the outer faces of the boundary tetrahedra. Using the definition of the basis functions, we obtain

$$b_i = -\frac{I}{A(\Gamma_2)} \sum_{e|i\in\partial\Omega^e, \partial\Omega^e\in\Gamma_2} \frac{A(\partial\Omega^e)}{3}, \tag{S.42}$$

where $A(\partial\Omega^e)$ is the area of the surface element $\partial\Omega^e$.

##### S.3.4.2 TMS

The TMS right-hand side is given by Equation S.23. Going through the same procedures as done with the stiffness matrix, we obtain

$$b_i = -\sum_{e|i\in\Omega^e} \int_{\Omega^e} \left(\underline{\sigma}\frac{\partial \boldsymbol{A}}{\partial t}\right) \cdot \boldsymbol{G}^e_{k^e_i}\, d\boldsymbol{x}. \tag{S.43}$$

Again, if we assume that $\left(\underline{\sigma}\frac{\partial \boldsymbol{A}}{\partial t}\right)$ is constant or varies linearly within each element, we can calculate the integrals exactly with only one integration point. Furthermore, results shown in the main paper text suggest that we can also use one-node integration in realistic applications of TMS simulations with only minor loss in accuracy.

#### S.3.5 Imposing Dirichlet Boundary Conditions

To impose Dirichlet boundary conditions of the form

$$\phi = a \quad \text{on } \Gamma \tag{S.44}$$

we use the elimination method. Suppose we have assembled the matrices, obtaining a linear system

$$\begin{bmatrix} S_{11} & \dots & S_{1i} & \dots & S_{1N} \\ \vdots & \ddots & \vdots & \ddots & \vdots \\ S_{i1} & \dots & S_{ii} & \dots & S_{iN} \\ \vdots & \ddots & \vdots & \ddots & \vdots \\ S_{N1} & \dots & S_{Ni} & \dots & S_{NN} \end{bmatrix} \begin{bmatrix} \phi_1 \\ \vdots \\ \phi_i \\ \vdots \\ \phi_N \end{bmatrix} = \begin{bmatrix} b_1 \\ \vdots \\ b_i \\ \vdots \\ b_N \end{bmatrix}, \tag{S.45}$$

and that we want to apply the boundary conditions to the node $\phi_i$. We begin by substituting the $i$-th row of the stiffness matrix by a row composed of zeroes if $i \neq j$ and 1 if $i = j$. We also replace the $i$-th entry of the RHS vector $\boldsymbol{b}$

$$\begin{bmatrix} S_{11} & \dots & S_{1i} & \dots & S_{1N} \\ \vdots & \ddots & \vdots & \ddots & \vdots \\ 0 & \dots & 1 & \dots & 0 \\ \vdots & \ddots & \vdots & \ddots & \vdots \\ S_{N1} & \dots & S_{Ni} & \dots & S_{NN} \end{bmatrix} \begin{bmatrix} \phi_1 \\ \vdots \\ \phi_i \\ \vdots \\ \phi_N \end{bmatrix} = \begin{bmatrix} b_1 \\ \vdots \\ a \\ \vdots \\ b_N \end{bmatrix}. \tag{S.46}$$

However, with these modifications the stiffness matrix loses its symmetry, which hampers the application of efficient sparse matrix solvers. We can recover the symmetry by the process of Gaussian elimination, multiplying each row by $S_j \leftarrow S_j - S_{ji}S_i$, we obtain

$$\begin{bmatrix} S_{11} & \dots & 0 & \dots & S_{1N} \\ \vdots & \ddots & \vdots & \ddots & \vdots \\ 0 & \dots & 1 & \dots & 0 \\ \vdots & \ddots & \vdots & \ddots & \vdots \\ S_{N1} & \dots & 0 & \dots & S_{NN} \end{bmatrix} \begin{bmatrix} \phi_1 \\ \vdots \\ \phi_i \\ \vdots \\ \phi_N \end{bmatrix} = \begin{bmatrix} b_1 - S_{ji}a \\ \vdots \\ 0 \\ \vdots \\ b_N - S_{ji}a \end{bmatrix}. \tag{S.47}$$

The system thus regains symmetry. We can remove the $i$-th row and column from the system, obtaining a modified system without $\phi_i$

$$\begin{bmatrix} S_{11} & \dots & S_{1N} \\ \vdots & \ddots & \vdots \\ S_{N1} & \dots & S_{NN} \end{bmatrix} \begin{bmatrix} \phi_1 \\ \vdots \\ \phi_N \end{bmatrix} = \begin{bmatrix} b_1 - S_{1j}a \\ \vdots \\ b_N - S_{Nj}a \end{bmatrix} \tag{S.48}$$

### S.4 Pairwise Analysis of Numerical Errors

To evaluate the numerical errors, we compared the meshes with nominal node densities of 0.125, 0.25, 0.5 and 1.0 nodes/mm$^2$ and their versions that have been refined by splitting. The number of nodes, elements and edge lengths for these models can be seen in Tables 2 and 3 of the main article. The results were compared both for the TMS and the TES setup shown in the main article, and the error evaluated using Equation 6 of the main article, with the electric field obtained with the high-resolution model as the reference electric field.

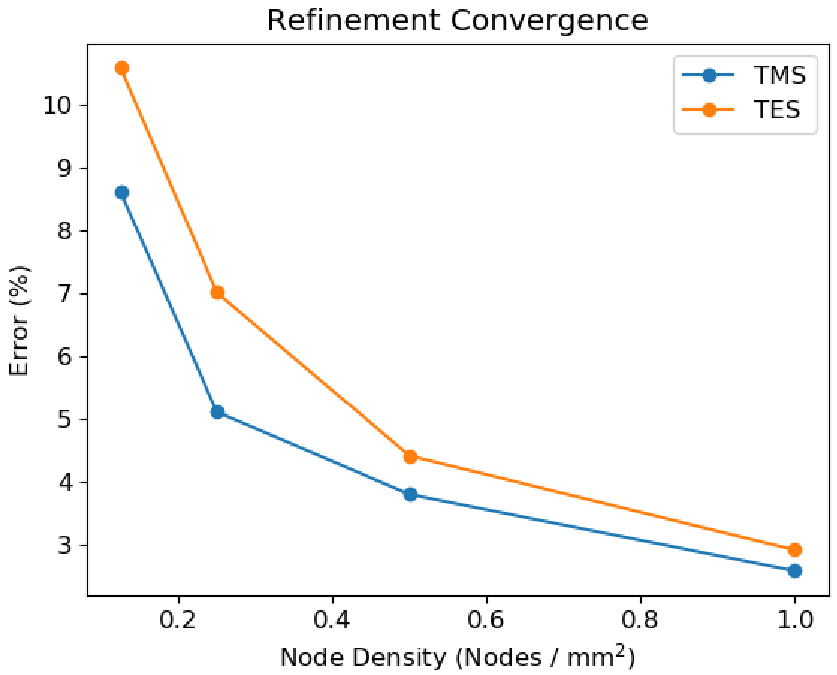


Figure S.1: Errors obtained for the TMS and TES electric fields when comparing the original and refined versions of the same head meshes

The results are shown in Figure S.1. We see that, as we improve the original head model, not only the *anatomical* accuracy (obtained by a better representation of the surfaces) of the solutions improved, but also the *numerical* accuracy (obtained by a larger functional space $V^h$), evidenced by the smaller error in relation to the refined mesh. However, as seen in Figure 8a of the main article, a better *numerical* accuracy does not necessarily translate into a better overall accuracy, when comparing with an anatomically more accurate head model.

[2] Pavel Ŝolín. *Partial differential equations and the finite element method*, volume 73. John Wiley & Sons, 2005.

[3] Weiping Wang and Solomon R Eisenberg. A three-dimensional finite element method for computing magnetically induced currents in tissues. *IEEE Transactions on Magnetics*, 30(6):5015–5023, 1994.

[4] J.Z Zienkiewicz, O.C. , Taylor, R.L, Zhu. *The Finite Element Method: its Basis and Fundamentals.* Elsevier, 2013.
